# Prior Scene Context Shapes the Neural Dynamics of Face Detection

**DOI:** 10.64898/2026.01.27.701765

**Authors:** Sule Tasliyurt-Celebi, Daniel Kaiser, Katharina Dobs

## Abstract

Detecting faces in our surroundings typically only takes a fraction of a second. How can such a rapid perceptual process still be influenced by prior expectations? Here, we used electroencephalography (EEG) to investigate how prior scene context modulates the temporal dynamics of neural face representations. Participants viewed natural scenes containing a single face (left or right), each preceded by either a faceless version of the same scene (preview condition) or a gray screen (no-preview condition), while performing a face detection task. Using multivariate decoding, we found that face location could be decoded shortly after target onset. Critically, decoding accuracy was higher in the preview condition at early stages, whereas the no-preview condition showed higher decoding at later time points, suggesting rapid facilitation by prior context followed by compensatory processing when contextual information was absent. In contrast, time-frequency decoding revealed a sustained preview advantage across alpha, beta, and gamma bands, even during time periods when evoked responses favored the no-preview condition. This dissociation between evoked and induced neural signals indicates that prior scene context engages distinct neural processes: early evoked activity reflects rapid contextual facilitation of sensory representations, while induced oscillatory activity may support prolonged context-dependent modulation. Together, these results show how prior scene context and sensory-driven processing jointly shape rapid face perception through temporally and spectrally distinct neural dynamics.

**Significance Statement:** Scientists studying perception have long investigated how the brain combines incoming sensory input with prior knowledge and expectations. Very fast perceptual decisions, however, are often assumed to rely primarily on sensory input alone. Here, we show that prior scene context shapes the neural processing of faces even during rapid face detection. When observers had prior information about a scene, neural representations of face location emerged earlier and followed a different temporal profile than when no contextual information was available. These findings demonstrate that even rapid face detection reflects an interaction between sensory-driven and context-dependent processes, and that prior knowledge influences perception through multiple, temporally distinct neural mechanisms.

## Introduction

Humans detect faces rapidly and accurately across diverse visual environments (Bindemann and Lewis, 2013). Face-directed saccades can occur as early as 100 ms after stimulus onset, faster than to any other object category (Crouzet et al., 2010; Martin et al., 2018), highlighting the remarkable efficiency of the visual system. This efficiency is often attributed to rapid bottom-up sensory processing, based on the speed and apparent automaticity of face detection. Moreover, faces reliably capture attention and are among the most frequently fixated objects in natural scenes (Fletcher-Watson et al., 2008; Cerf et al., 2009). Such rapid orienting toward faces has been interpreted as evidence that face detection is largely stimulus-driven and minimally influenced by top-down factors such as task goals or scene context (Crouzet and Thorpe, 2011). Consistent with this view, MEG signals indicate the presence of faces within the first 100 ms of processing (Cauchoix et al., 2014), and M/EEG decoding studies show that face category and identity can be reliably distinguished within 150-200 ms after stimulus onset (Ambrus et al., 2019; Dobs et al., 2019). Together, these findings suggest that the visual system rapidly transforms sensory input into informative face representations.

The fast and seemingly automatic nature of face detection raises the question of to what extent this process can nevertheless be modulated by top-down factors. Accumulating evidence indicates that even rapid perceptual decisions can be shaped by expectations, prior knowledge, and task demands (Gregory, 1970; Summerfield and Egner, 2009). In line with this view, recent behavioral studies show that prior information and contextual expectations can facilitate face perception (Garlichs and Blank, 2024; Mares et al., 2024; Tasliyurt-Celebi et al., 2024). In our work, prior information refers to expectations derived from the spatial layout of a scene before a face appears. We previously showed that prior scene previews enhance face detection from the very first saccade onward, and that the degree of mismatch between the predicted location of a face in a scene and its actual position systematically predicts face detection latency (Tasliyurt-Celebi et al., 2024). However, the neural mechanisms underlying such rapid context-dependent modulation remain unclear.

Previous work suggests that expectations bias perceptual processing by modulating neural representations of visual information. Multivariate decoding studies have shown that expectations alter how simple visual features are encoded in neural activity (de Lange et al., 2018), and object-based attention enhances representations of task-relevant categories as early as 150–200 ms after stimulus onset (Kaiser et al., 2016). In addition to these temporal effects, expectations also influence rhythmic neural activity. Low-frequency oscillations in the alpha and beta range have been linked to top-down processes such as expectation, attention, and predictive feedback, often showing enhanced representations of predictable stimuli (Noah et al., 2020; Chen et al., 2023; Stecher et al., 2025). However, whether such expectation-related mechanisms also modulate the rapidly emerging, seemingly bottom-up driven neural responses that support face detection remains an open question.

Here, we combine multivariate pattern decoding with high-temporal-resolution EEG to characterize the neural mechanisms underlying rapid face detection. In a face localization task, we first replicated that providing a brief, faceless preview of the scene facilitates face localization: participants were faster to indicate whether a face appeared on the left or right when they had seen the scene beforehand. Building on this behavioral effect, we recorded EEG activity while participants performed a face present/absent detection task with and without prior scene preview. Crucially, face location was decoded from neural activity while remaining orthogonal to task demands. Using time-resolved and time-frequency-resolved decoding analyses, we investigated when and in which frequency bands information about face location emerged. This approach allowed us to test how prior scene context modulates early sensory representations as well as more sustained oscillatory activity, thereby providing insight into how contextual predictions interact with rapid visual processing in the human brain.

## Method

### Behavioral Face Localization Task

#### Participants

Twenty-four participants (5 male; M = 23.87 years, SD = 4.72) with normal or corrected-to-normal vision were recruited from the university mailing list of the Justus Liebig University Giessen (JLU). Participants received either €8 or one course credit as compensation and provided written informed consent prior to participation. The study was approved by the local ethics committee of JLU and conducted in accordance with the Declaration of Helsinki.

#### Stimuli

To test the role of prior scene information on face location in natural scenes, we selected 784 images, each containing a single face. In half of the images (392), the face was positioned on the left side of the scene, and in the other half on the right. Most images were drawn from the Places365 dataset (Zhou et al., 2018), with additional images collected from the internet to balance the number of left- and right-sided stimuli. These images served as target stimuli. For each target image, we created a corresponding faceless version by manually editing out the person using Adobe Photoshop version 2022 (Adobe Inc., 2019). These images served as scene previews. To eliminate potential confounds related to face location within the scene, all stimuli (preview and target) were horizontally mirrored, and the assignment of mirrored versions was counterbalanced across participants. All stimuli were resized to a resolution of 800 × 600 pixels.

#### Experimental Design

To investigate how prior scene information influences face localization in natural scenes, participants performed a two-alternative forced-choice task indicating the side (left vs. right) on which a face appeared. Each trial began with the presentation of either a scene preview (faceless version; preview condition) or a gray screen (no-preview condition) for 250 ms. This was followed by a static noise pattern displayed for 50 ms and a fixation cross for 500 ms to minimize residual visual traces of the preview and to avoid a “face pop-out” effect when the subsequent target appeared. The target stimulus was then presented for 100 ms, and participants were instructed to press the corresponding left or right arrow key as fast as possible to indicate face location.

Each participant completed 784 trials in total, equally divided between the preview and no-preview conditions (392 trials each). Mirrored versions of all stimuli were used to ensure balance between left- and right-sided faces, thereby eliminating potential scene-related biases. The full stimulus set was split into four subsets and counterbalanced across participants such that each image appeared equally often in all conditions (original/mirrored x preview/no-preview). Trials were divided into eight blocks of 98 stimuli each, with short breaks between blocks. Each block lasted approximately five minutes. Trial order was randomized for each participant. Before the main experiment, participants completed 10 practice trials to familiarize themselves with the task.

#### Statistical analysis

To compare behavioral measures (e.g., reaction time, accuracy) between preview and no-preview conditions, we used paired *t*-tests. Pearson correlations were used when assessing relationships within the same dataset (e.g., split-half reliability). Statistical significance was evaluated at *p* < .05 (two-tailed), and effect sizes (Cohen’s *d*) are reported where appropriate.

### EEG Face Detection Task

#### Participants

Forty-four participants (17 male; M = 26.27 years, SD = 4.31) with normal or corrected-to-normal vision took part in the EEG study. Data from one participant were excluded due to poor signal quality resulting from persistently high electrode impedances. All participants provided written informed consent prior to participation and received either one course credit or €12 per hour as compensation. The study was approved by the local ethics committee of JLU and conducted in accordance with the Declaration of Helsinki.

### Stimuli

To examine the influence of prior scene information on neural face representations during face detection, we used the same stimulus set (784 images, each containing a single face) as in the behavioral face localization experiment. To increase the number of unique trials in the EEG study, participants viewed both the original and mirrored versions of each stimulus. One version was presented in the preview condition and the other in the no-preview condition, counterbalanced across participants to avoid stimulus repetition within a condition. Additionally, 100 images without faces served as foil trials, each with a mirrored counterpart.

#### Experimental Design

To avoid confounding neural representation of face location with overt localization responses, participants performed an orthogonal face detection task during the EEG experiment. The procedure was identical to that of the behavioral face localization experiment up to the target presentation. The target stimulus was presented for 100 ms, and participants were instructed to decide whether the scene contained a face. If no face was present, participants pressed the spacebar; if a face was detected, no response was required. Responses could be made from target onset until the beginning of the next trial (1200 ms in total). A fixation cross remained visible throughout the experiment to promote sustained attention and minimize eye and head movements.

The stimulus set was divided into two halves (part 1 and part 2). Assignment of these stimulus parts to conditions was counterbalanced across participants: half of the participants saw part 1 in the preview condition and part 2 in the no-preview condition, while the remaining participants saw the opposite pairing. To maximize the number of trials, each participant viewed both the original and mirrored versions of each stimulus; however, each version appeared in a different condition (i.e., one with preview, the other without). In total, participants completed 1,768 trials (784 original and 784 mirrored face-containing stimuli), including 200 foil trials (100 original and 100 mirrored), evenly distributed across preview and no-preview conditions. Each version (original or mirrored) was shown once per participant and was preceded either by a scene preview or a gray screen. Trial assignment (original/mirrored x preview/no-preview) was counterbalanced across participants, and trial order was randomized. To assess whether presenting original and mirrored versions introduced familiarity effects, we performed a control analysis comparing decoding for the first versus second presentation (Supplementary Note 1; Supplementary Fig. S1).

EEG was recorded simultaneously while participants performed the face detection task. The experiment consisted of 13 blocks of 136 trials each, separated by short breaks. During each break, the 9-point calibration and validation procedure was repeated to maintain recording accuracy. Prior to the main experiment, participants completed 10 practice trials to familiarize themselves with the task. Including preparation and setup, the entire session lasted approximately 2.5 hours. Participants performed the face-detection task with high accuracy (87%). Performance did not differ between preview and no-preview conditions (paired t test; p = .51).

### EEG data acquisition and preprocessing

Participants were seated comfortably in a dimmed room at a fixed distance of 57 cm from the monitor, using a combined forehead-and-chin rest to minimize head movement. EEG was recorded with a 64-channel Easycap system (Brain Products GmbH) at a sample rate of 1,000 Hz. Fz served as the reference electrode and AFz as the ground. Electrode impedances were kept below 20 kΩ, and electrodes were arranged according to the international 10–10 system. Event triggers were transmitted via a parallel port from the presentation computer to the EEG acquisition system.

Continuous EEG data were bandpass filtered between 0.1 and 100 Hz. Noisy channels (M = 2.77 ± 2.72) were identified by visual inspection and replaced with the average of their neighboring electrodes. Independent component analysis (ICA) was performed to remove eye movement artifacts, with components identified through visual inspection based on characteristic scalp topographies and time courses. The cleaned data were segmented into epochs between -100 ms and 1900 ms relative to target onset and baseline-corrected using the -100 to 0 ms prestimulus interval. Data were then downsampled to 200 Hz. Foil trials were excluded from all analyses. Preprocessing was conducted using the MNE toolbox in Python (Gramfort et al., 2013).

#### Decoding Analysis

We applied multivariate pattern analysis (MVPA) using a linear support vector machine (SVM) classifier to decode the spatial position of faces (left vs. right) from EEG data at the participant level. To improve signal-to-noise ratio, pseudo-trials were constructed at each time point by averaging the data within the same face-location (left vs. right) and condition (preview vs. no-preview) into 10 folds (Fig. 3a). For evoked responses, each feature pattern consisted of the sensor activations corresponding to one pseudo-trial and condition. Decoding was performed in a time-resolved manner using a sliding window of 50 ms width, advanced in 5 ms steps from –100 to 1100 ms relative to target onset. Classifiers were trained on nine folds and tested on the remaining fold, and decoding accuracies were averaged across all cross-validation splits. At the group level, decoding accuracy was tested against chance level using a sign-permutation test with cluster-based correction across time (5,000 iterations); (Maris and Oostenveld, 2007; Gramfort et al., 2013), controlling for multiple comparisons over time. All analyses were conducted using the MNE toolbox in Python (Maris and Oostenveld, 2007; Gramfort et al., 2013).

As a control, we repeated the decoding procedure during the preview interval to test whether eventual face location was decodable from preview-evoked activity (Supplementary Note 2; Supplementary Fig. S2).

### Time-Frequency Analysis

Time-frequency analysis was performed using a Morlet wavelet transformation (6-cycle length) to extract oscillatory activity in the theta (4–7 Hz), alpha (8–13 Hz), beta (14–30 Hz) and lower gamma (31–40 Hz) frequency bands over the interval from –100 to 1100 ms relative to target onset. Frequency-resolved data were then averaged across all frequencies within each band. For each frequency band, we performed the same decoding analysis as for the evoked responses (see Decoding Analysis), training linear SVM classifiers to decode face location (left vs. right) separately for each condition.

## Results

### Prior scene information enhances face detection and spatial localization

Before the EEG experiment, we conducted a behavioral study to test whether prior scene information facilitates face localization. Participants (N = 24) performed a face localization task in which they indicated whether a briefly presented face appeared on the left or right side of a natural scene. Mean response time across conditions was 464 ms. Critically, participants localized faces significantly faster in the preview condition than in the no-preview condition (Fig. 2; 451 ms vs. 476 ms, *t*(23) = 5.42, *p* = .000, Cohen’s d = .27), indicating that prior scene information facilitates spatial localization of faces.

**Fig. 1.**
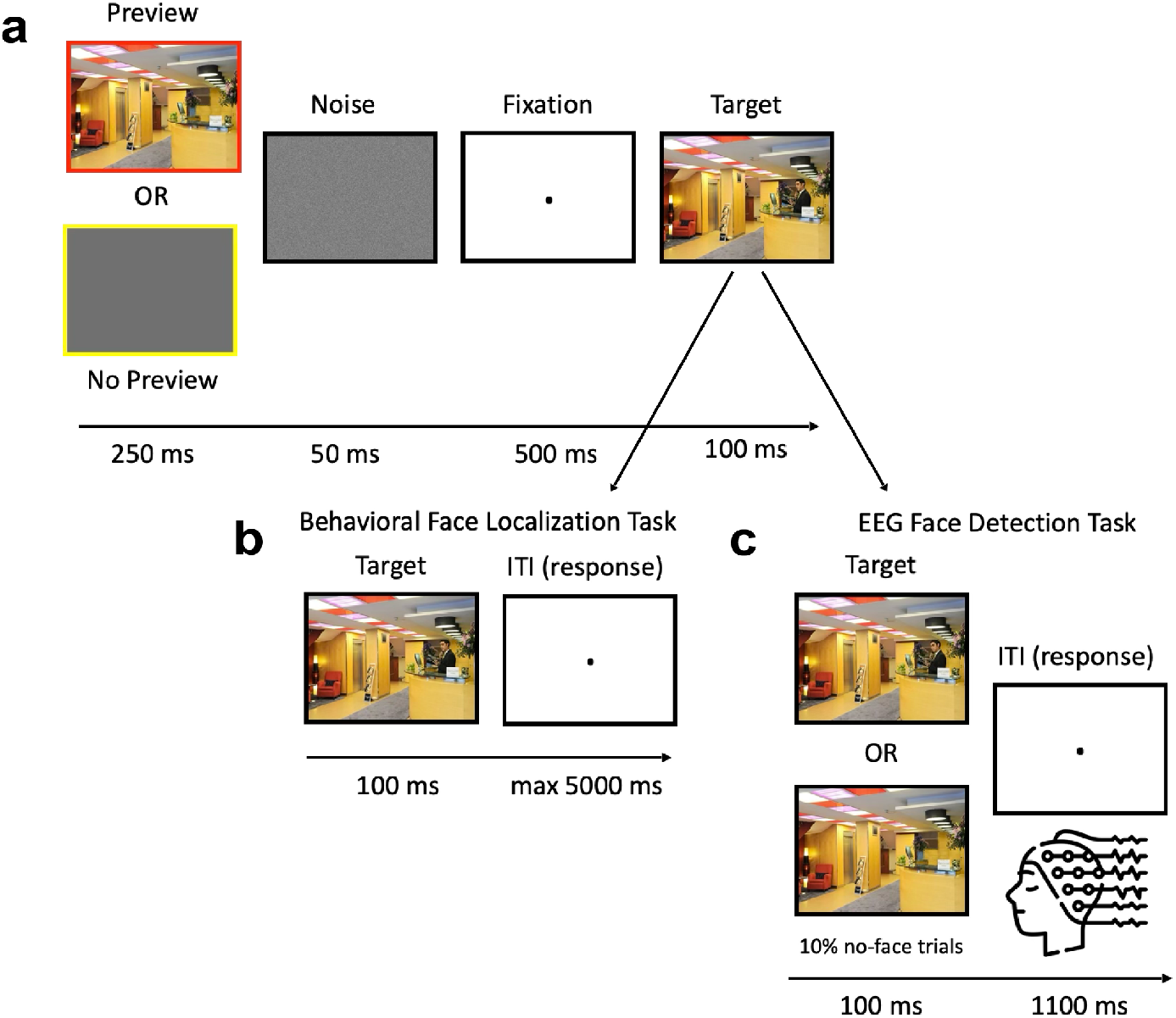
Procedure for both tasks. **a**, Participants were shown either a faceless preview (preview condition) or a gray screen (no-preview condition) before viewing the target scene. **b**, Behavioral face localization task. Participants (N=24) decided where the face was located (left or right) in the scene. **c**, EEG face detection task. Participants (N=44) were instructed to decide whether there was a face in the target scene and press a button if there was none. All scene images were taken from the Places365 database and are used for non-commercial research purposes. These images do not raise any conflict with the copyright policy of the database.

**Fig. 2.**
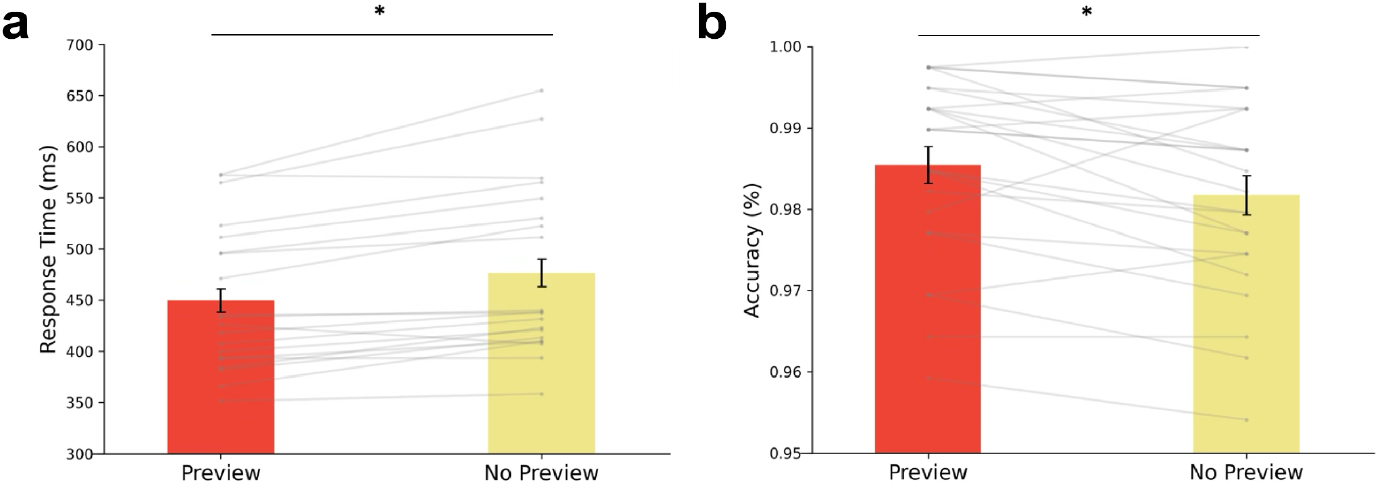
Behavioral results of the face localization task. **a**, Response times were significantly shorter in the preview than in the no-preview condition. **b**, Localization accuracy was significantly higher in the preview than in the no-preview condition. Gray lines represent individual participants. Error bars denote ±SEM. Asterisks indicate *p* < .01 (paired t-test).

**Fig. 3.**
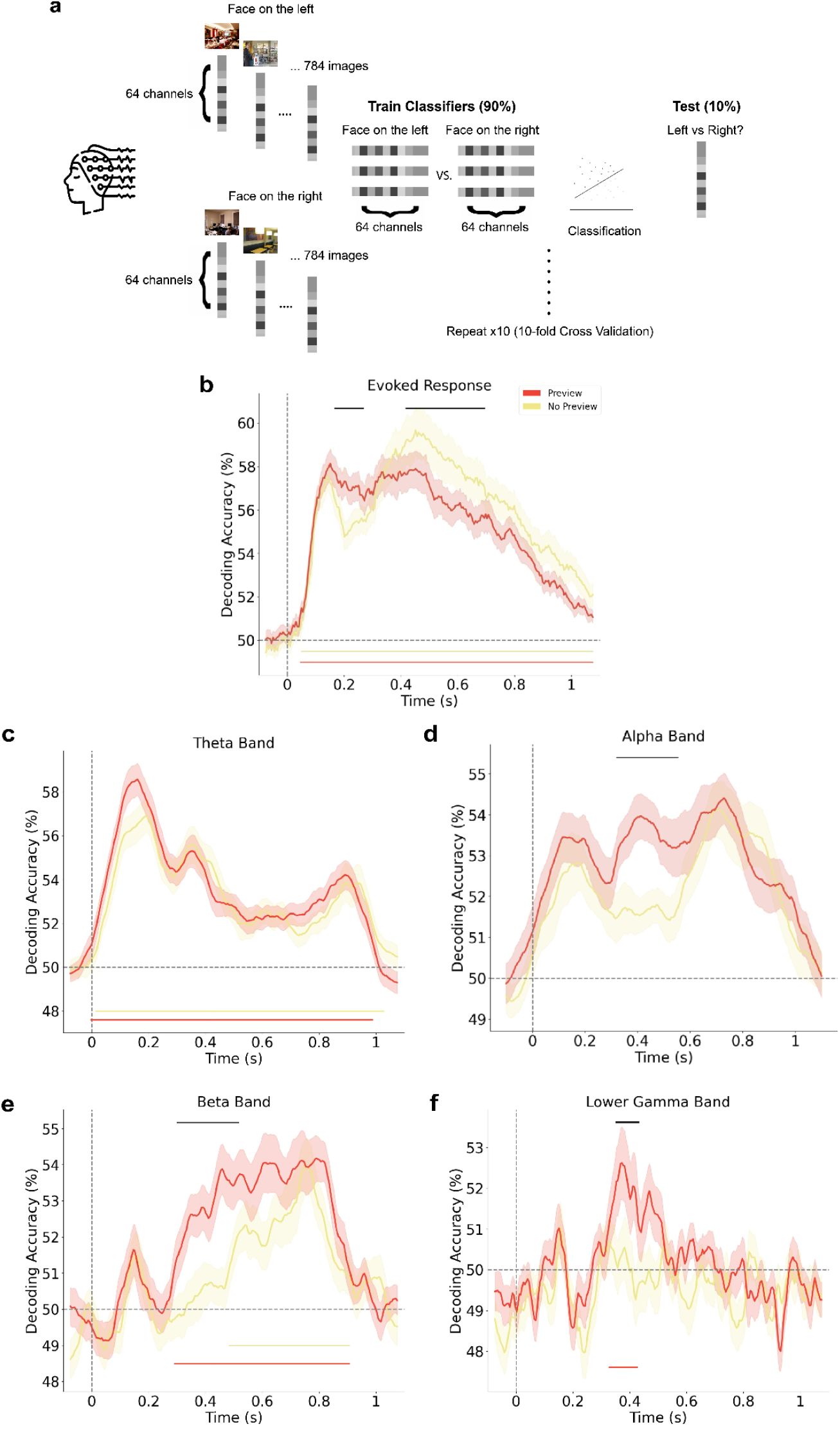
EEG decoding procedure and results. **a**, Schematic of the decoding analysis. Linear SVM classifiers were trained within each condition (preview vs. no-preview) to discriminate face location (left vs. right) using 10-fold cross-validation. Decoding was applied to both time-locked (broadband) and time-frequency-resolved EEG data. **b**, Time-resolved decoding accuracy from evoked (broadband) responses. Shaded areas represent ± SEM across participants (N = 43). **c–f**, Time-frequency-resolved decoding accuracy for the theta (4-7 Hz), alpha (8–13 Hz), beta (14–30 Hz), and lower gamma (31–40 Hz) bands. Red indicates preview condition, yellow no-preview condition. Colored horizontal lines indicate time intervals with decoding accuracy significantly above chance (*p* < 0.05, sign permutation test, cluster-corrected). Black horizontal lines indicate time windows with significant differences between conditions (*p* < 0.05, cluster-corrected). All scene images were taken from the Places365 database and are used for non-commercial research purposes. These images do not raise any conflict with the copyright policy of the database.

Localization accuracy was high overall and showed a small but significant increase in the preview condition (preview: *M* = 98.54%, *SD* = 0.01; no-preview: *M* = 98.17%, *SD* = 0.01; *t*(23) = 2.92, *p* = .008, Cohen’s d = .32), ruling out a speed-accuracy trade-off.

To assess image-level reliability, we computed split-half correlations of response times within each condition, revealing moderate consistency (preview: *r* = .58; no-preview: *r* = .49; Spearman–Brown corrected). To test whether the behavioral preview benefit was image-specific, we computed the preview effect (reaction time difference between no-preview and preview conditions) for each image and assessed its reliability across split halves of participants. This analysis revealed low reliability of the image-wise preview effect (r = .19), indicating that the magnitude of preview-related facilitation was not consistently tied to specific images. In contrast, overall image-wise response times collapsed across conditions showed high reliability (r = .75), demonstrating robust and consistent stimulus-specific difficulty.

Together, these results suggest that prior scene previews exert a global facilitative effect on face localization rather than selectively benefiting a stable subset of images.

### Prior scene information facilitates early neural encoding of face location

Having established that prior scene context enhances face localization at the behavioral level, we next asked how this facilitation is implemented in the brain. Specifically, we asked whether prior scene context modulates neural representations of face location during detection, and which mechanisms mediate the interaction between bottom-up sensory processing and top-down contextual influences. To address these questions, we used a very closely matched experimental paradigm while recording EEG. In contrast to the behavioral face localization task, participants (N = 43) performed a face detection task that was orthogonal to the decoded dimension (face location), allowing us to probe neural representations of spatial face information independent of task demands. We used multivariate decoding to characterize the temporal dynamics of face-related neural signals.

We were able to decode face location from evoked EEG responses as early as ∼46 ms after target onset, and decoding remained above chance throughout the trial (preview: max. accuracy = 58.1% at 151 ms; no-preview: max. accuracy = 59.7% at 452 ms; Fig. 3b). Critically, decoding accuracy was higher in the preview condition at early processing stages (166–266 ms, maximum *d* = .56), whereas the no-preview condition showed greater decoding at later stages (417–693 ms, maximum *d* = .69 ; Fig 3b). This temporal dissociation suggests an early facilitation of neural face-location representations by prior scene context, followed by later compensatory processing when such context is absent.

To complement the time-locked decoding analyses and to probe frequency-specific mechanisms potentially involved in contextual modulation, we next examined oscillatory activity in the EEG signal. We trained and tested linear classifiers on time-frequency-resolved data separately for the theta (4-7 Hz), alpha (8–13 Hz), beta (14–30 Hz), and lower gamma (31-40 Hz) bands to discriminate face location (left vs. right).

Decoding analyses revealed significantly higher decoding accuracy in the preview condition in the alpha band (191–347 ms, maximum *d* = .46, and 362-573 ms, maximum *d* = .64; Fig. 3c), the beta band (300–517 ms; maximum *d* = -.31; Fig. 3d), and the lower gamma band (268–402 ms, maximum *d* = -.33; Fig. 3e). Enhanced decoding in the alpha and beta bands is consistent with prior work linking these frequencies to top-down spatial orienting and predictive influences (Thut et al., 2006; van Kerkoerle et al., 2014; Michalareas et al., 2016; Bastos et al., 2020; Stecher et al., 2025), whereas increased decoding in the lower gamma range may be consistent with facilitated feedforward processing of task-relevant visual information (Fries, 2009; Kuzovkin et al., 2018; Brunet and Fries, 2019). Notably, for all three frequency bands, the preview advantage persisted during time windows in which time-locked broadband decoding showed a relative advantage for the no-preview condition. Thus, induced oscillatory activity continued to reflect enhanced face-location information when prior scene context was available, even as evoked responses suggested compensatory processing in its absence. In contrast, decoding in the theta band showed no reliable difference between conditions at any time point, suggesting that theta-range activity did not contribute to the contextual modulation. Together, these findings reveal a dissociation between time-locked broadband responses and induced rhythmic activity, suggesting that prior scene context engages distinct neural mechanisms that unfold across both time and frequency domain.

## Discussion

The goal of this study was to characterize how prior scene context modulates the neural dynamics of rapid face detection. We found that scene previews facilitated both behavioral performance and neural representations of face location in natural scenes. Behaviorally, participants localized faces faster and more accurately when a scene preview was available, suggesting that prior contextual information supports more efficient orienting toward likely face locations. This aligns with our previous work showing that scene previews facilitate saccadic targeting toward faces (Tasliyurt-Celebi et al., 2024). At the neural level, multivariate decoding revealed enhanced face-location information shortly after target onset in the preview condition, consistent with recent evidence that expectations and context can influence early stages of face processing (Garlichs and Blank, 2024; Mares et al., 2024). In contrast, face-location decoding was stronger at later stages in the no-preview condition, suggesting prolonged stimulus-driven processing or later-stage decision-related accumulation when contextual information is unavailable. Crucially, time-frequency decoding showed a sustained preview advantage across alpha, beta, and gamma bands, whereas no comparable enhancement for the no-preview condition was observed in induced oscillatory signals. Together, our results suggest that prior scene context enhances early evoked face-location representations and sustained induced signals, whereas the absence of context elicits delayed, compensatory time-locked decoding.

The timing of the decoding effects offers important insight into how early contextual expectations interact with rapid face processing. Although faces can elicit extremely fast behavioral and neural signatures (Cauchoix et al., 2014; Martin et al., 2018; Dobs et al., 2019), the preview-related early enhancement we observe emerges in a time window (∼170-290 ms) that is not substantially earlier than top-down modulations reported for other types of visual processing. For example, object-based attention and expectation manipulations have been shown to modulate multivariate stimulus representations at ∼150–200 ms post-stimulus (Kaiser et al., 2016). The similarity in timing suggests that, even for faces, the influence of prior knowledge may rely on domain-general mechanisms that interact with perceptual processing at mid-latency stages, rather than reflecting a uniquely accelerated “shortcut” for faces. The advantage for faces over other categories may derive from a more efficient readout of these amplified representations during downstream processing. Importantly, this temporal profile is consistent with an account in which contextual information biases early perceptual encoding during online processing (e.g., by sharpening or prioritizing spatially relevant signals), rather than reflecting a late post-perceptual stage. Conversely, the later enhancement of face-location decoding in the no-preview condition suggests that when contextual information is unavailable, the system may require a more prolonged stimulus-driven analysis to infer face location, resulting in delayed compensatory representations. Alternatively, this delayed enhancement could reflect increased uncertainty and weaker contextual priors, leading the system to weight stimulus-driven evidence more strongly at later processing stages.

Enhanced decodability in the alpha and beta bands between 200-600 ms suggests that prior scene context modulates face processing through frequency channels commonly linked to top-down influences, such as expectation and spatial orienting (Stecher et al., 2025). Alpha-band oscillations, in particular, have been repeatedly associated with spatial attention and the prioritisation of relevant locations in visual space (Samaha et al., 2018), whereas alpha/beta rhythms more broadly have been proposed to carry predictive or feedback-related signals in hierarchical vision (van Kerkoerle et al., 2014; Michalareas et al., 2016). In the present paradigm, the observed alpha/beta preview advantage could reflect (at least) two non-exclusive mechanisms. First, alpha may reflect a spatial orienting process triggered by contextual cues, which would support more efficient sampling or prioritisation of the expected face location. Importantly, given that the alpha/beta effect emerges after the earliest enhancement in evoked decoding, this orienting signal may not purely reflect a preparatory pre-target bias, but could instead reflect post-target attentional orienting toward the detected face location, consistent with object-based attentional selection constraining spatial orienting in real-world scenes (Malcolm and Shomstein, 2015). Second, the alpha/beta effects may reflect sustained feedback-driven modulation of sensory processing, as proposed by predictive routing frameworks in which top-down rhythms gate or modulate feedforward information flow (Bastos et al., 2015). This interpretation is also consistent with the fact that the preview advantage in frequency-based decoding persists during time windows where evoked responses show compensatory processing in the no-preview condition, suggesting that induced oscillatory signals capture sustained context-related modulation that is not time-locked to the sensory transient. Notably, the presence of a preview advantage in lower gamma may reflect facilitated feedforward transmission under contextual guidance, potentially via coordinated interactions between alpha/beta feedback and gamma feedforward channels (Bastos et al., 2015; Fries, 2015). Together, these findings suggest that prior scene context engages temporally and spectrally distinct signatures of contextual modulation, likely indicating that prior experience modulates the sustained interplay of context-dependent and sensory-driven processes that follows on from the initial facilitation effects observed in evoked broadband signals.

Our findings raise the question of which computational mechanisms could implement such context-sensitive neural dynamics during rapid face perception. Many state-of-the-art computational models of face perception, particularly feedforward deep neural networks, lack explicit mechanisms for integrating prior context or top-down influences (O’Toole and Castillo, 2021; van Dyck and Gruber, 2023). However, contextual facilitation does not necessarily require anatomical recurrence or feedback: in principle, prior information could bias processing in an otherwise feedforward hierarchy, for example via rapid gain modulation or pre-activation of feature channels relevant for the expected target (Lindsay and Miller, 2018). At the same time, a second possibility is that contextual effects reflect recurrent or feedback computations, consistent with predictive-coding accounts in which priors are integrated with sensory evidence through iterative inference (Egner et al., 2010; Spratling, 2017; Islah et al., 2025). This opens a promising modeling avenue: Contrasting biased feedforward networks with recurrent networks (Kar et al., 2019; Kietzmann et al., 2019) could determine which mechanisms best capture the temporal dissociation observed in the EEG data (rapid ERP facilitation vs. sustained oscillatory enhancement). Such comparisons may help clarify whether the preview effect arises from changes in the pre-stimulus neural state that bias subsequent feedforward processing, from recurrent or feedback integration, or both.

While our study provides converging behavioral and neural evidence for context facilitation of rapid face detection, several open questions remain. First, our manipulation used an identical faceless preview of the upcoming scene, which cleanly isolates prior scene information but leaves open how expectations operate in more variable natural settings (Võ and Wolfe, 2012). Future studies could test whether the observed facilitation generalizes to less stimulus-specific previews, for example, different examples from the same scene category, or previews that preserve only coarse layout statistics. This would also help disentangle stimulus-specific priming/preactivation (e.g., low-level layout or mid-level structure) from more abstract category-based expectations. Second, because faces in natural scenes are often accompanied by bodies, part of the preview benefit may reflect facilitation of person-related cues more generally rather than faces alone. Future studies could dissociate face and body contributions by independently manipulating their visibility (Hickey et al., 2015). Third, we decoded face location while participants performed an orthogonal detection task; this helps dissociate decoded content from task demands, but limits direct linkage to spatial behavior and calls for future work connecting neural dynamics to trial-wise behavior and eye movements.

In sum, our findings show that prior scene context modulates rapid face detection by reshaping the neural dynamics of face location representations. Scene previews enhance early time-locked decoding and induce sustained oscillatory representations across frequency bands, whereas the absence of context yields delayed compensatory decoding. These results provide neurophysiological constraints on predictive processing accounts, suggesting that contextual priors and sensory evidence interact through temporally and spectrally distinct mechanisms during naturalistic perception. More broadly, they highlight that even rapid visual decisions are not purely stimulus-driven, but emerge from dynamic integration of feedforward signals and contextual factors.

## Acknowledgments

K.D. was supported by the ERC Starting Grant DEEPFUNC (ERC-2023-STG-101117441). D.K. was supported by the German Research Foundation (DFG; KA4683/5-1, project number 518483074) and the ERC Starting Grant PEP (ERC-2022-STG-101076057). K.D. and D.K. were both supported by the DFG under Germany’s Excellence Strategy (EXC 3066/1 “The Adaptive Mind”, project number 533717223) and the Collaborative Research Center SFB/TRR 135 (project number 222641018). Views and opinions expressed are those of the authors only and do not necessarily reflect those of the funders. Neither the funders nor the granting authority can be held responsible for them.

## Data and code availability

All stimuli are available on OSF (https://doi.org/10.17605/OSF.IO/7WXQ4). All data and Python code (version 3.8.20) used for data preprocessing, analyses, and visualization will be made publicly available on GitHub (https://github.com/VCCN-lab) upon publication.

