## Supplementary Note 1 and 2 for "Prior Scene Context Shapes the Neural Dynamics of Face Detection"

### Supplementary Information

#### Supplementary Note 1

To assess whether stimulus familiarity due to mirroring of stimuli influenced decoding, we conducted a control analysis comparing neural decoding for the first versus second presentation of each image. In our design, each scene was presented twice—once in its original orientation and once mirrored—with the two versions assigned to different conditions (preview vs. no-preview; counterbalanced across participants). Although mirroring alters the spatial layout, participants may still recognize a scene upon its second encounter. If so, repetition-related effects could, in principle, affect decoding independently of the preview manipulation.

We therefore labeled trials as first or second occurrences and repeated the time-resolved decoding of face location using the same preprocessing and analysis pipeline as in the main experiment. As shown in Supplementary Figure S1, decoding time courses were highly similar for first and second occurrences, and no reliable differences were observed at any time point ($p > .05$). This suggests that repetition did not measurably modulate face-location decoding, supporting the interpretation that the main effects reflect modulation by prior scene context rather than repetition-related neural changes.

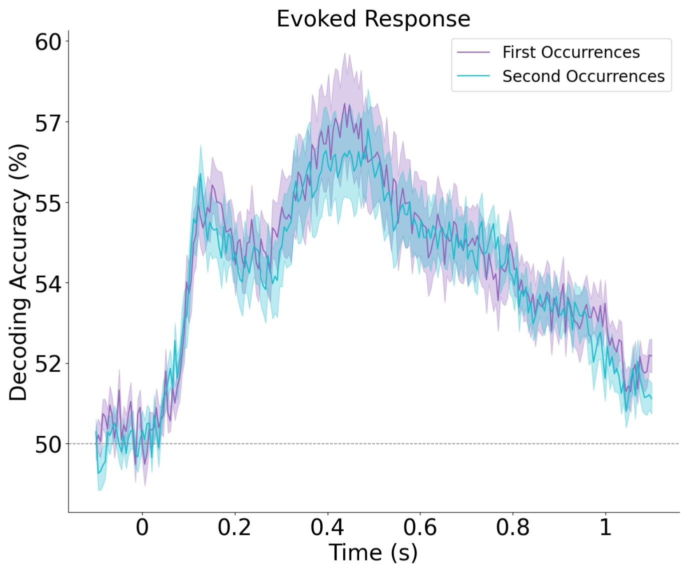


**Supplementary Fig. S1 |** Time resolved decoding accuracy for face location is shown separately for the first (purple) and second (blue) presentation of each scene. Each image was presented twice, once in its original and once mirrored version with presentations assigned to the different experimental conditions. Decoding accuracy rose above chance shortly after stimulus onset and followed highly similar time courses for first and second occurrences. No significant differences were observed at any time point (all $p > .05$). Shaded areas indicating SEM across participants.

#### Supplementary Note 2

To determine whether the preview display contained any lateralized neural information that could contribute to differences in face-location decoding, we conducted an additional decoding analysis on EEG activity during the preview interval. In our design, participants viewed either a faceless scene preview (preview condition) or a gray screen (no-preview condition) for 250 ms before target onset. The preview scene was always the faceless version of the upcoming target and, critically, did not contain cues indicating whether the face in the subsequent target would appear on the left or right. Nevertheless, we considered the possibility that the preview image might induce subtle spatial biases or anticipatory neural patterns that could influence later decoding.

To evaluate this possibility, we applied the same time-resolved multivariate decoding analysis used in the eeg experiment to the preview interval. EEG data were baseline-corrected using a −100 to 0 ms window relative to preview onset, and pseudo-trials were constructed in the same way as for the target decoding. We then trained a classifier to decode the eventual face location (left vs. right) based solely on neural activity during the preview display. This analysis was conducted only for the preview conditions. As shown in Supplementary Figure S2, decoding accuracy during the preview period remained at chance throughout the entire 250 ms interval, with no significant clusters observed ($p$ = 1). Thus, we found no evidence that face location could be predicted from preview-evoked EEG activity.

Together, these results provide no evidence that the preview interval carries decodable information about the upcoming face location. This suggests that the main decoding effects are not driven by preview-evoked lateralized activity. Notably, while the ability to predict face location from faceless scene structure would be theoretically interesting as a scene-based prior; our analysis indicates that such information is not reliably present (or not decodable with our methods) during the preview display.

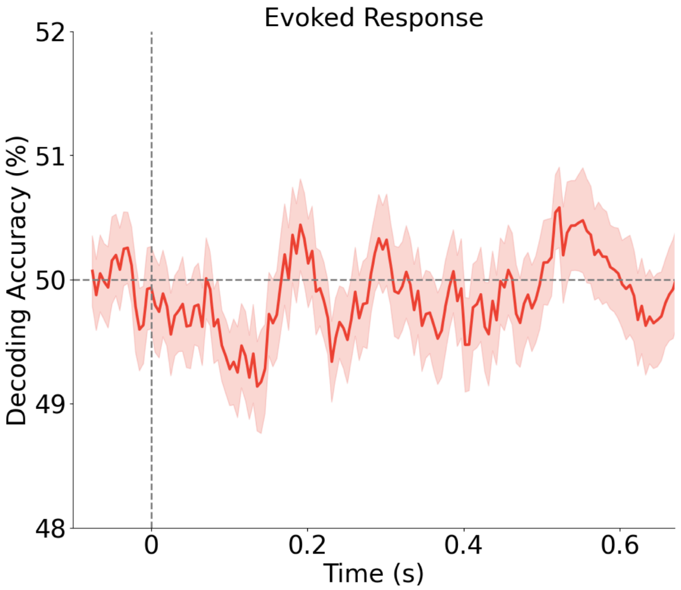


**Supplementary Fig. S2 |** Time-resolved decoding accuracy for *eventual face location* (left vs. right) during the preview interval, time-locked to preview onset (preview condition only). The dashed line denotes chance level.
